# Abnormal LTR retrotransposons formed by the recombination outside the host genome

**DOI:** 10.1101/452573

**Authors:** Fanchun Zeng, Zhongyi Sun

## Abstract

Long terminal repeat (LTR) retrotransposons are the dominant feature of higher plant genomes, which have a similar life cycle with retrovirus. Previous studies cannot account for all observed complex LTR retrotransposon patterns. In this study, we first identified 63 complex LTR retrotransposons in rice genome, and most of complex elements harbored flanking target-site duplications (TSDs). But these complex elements in which outermost LTRs had not the most highly homologous can’t be explained. We propose a new model that the homologous recombination of two new different normal LTR retrotransposon elements in the same family can occur before their integration to the rice genome. The model can explain at least fourteen complex retrotransposons formations. We also find that normal LTR retrotransposons can swap their LTRs to generate abnormal LTR retrotransposons in which LTRs are different because of homologous recombination before their integration to the genome.

## Background

Long terminal repeat (LTR) retrotransposons are the main composition of higher plant genomes, which are a major reason for plant genome amplification and shrinkage (Kumar and Bennetzen 1999; Dupeyron *et al*. 2017). The insert/delete activity of LTR retrotransposons is key for genome size change and evolution in plant. LTR retrotransposons have a similar process of replication to retrovirus (Sabot and Schulman 2006; Schulman 2013) that they are replicated in a “copy and paste” mode via RNA intermediates (Finnegan 1989) (Figure 1). The two terminals are LTRs that with an internal region in the middle, which may or not contains open reading frame (ORF). Insertion of an LTR retrotransposon to the host genome can form a new copy which harbors flanking target-site duplications (TSDs). The complex LTR retrotransposons which contain three LTRs (LTR1—LTR2—LTR3) had been identified in many species (Devos *et al*. 2002; Sabot and Schulman 2007), but their formation mechanism was still unclear.

**Figure 1:**
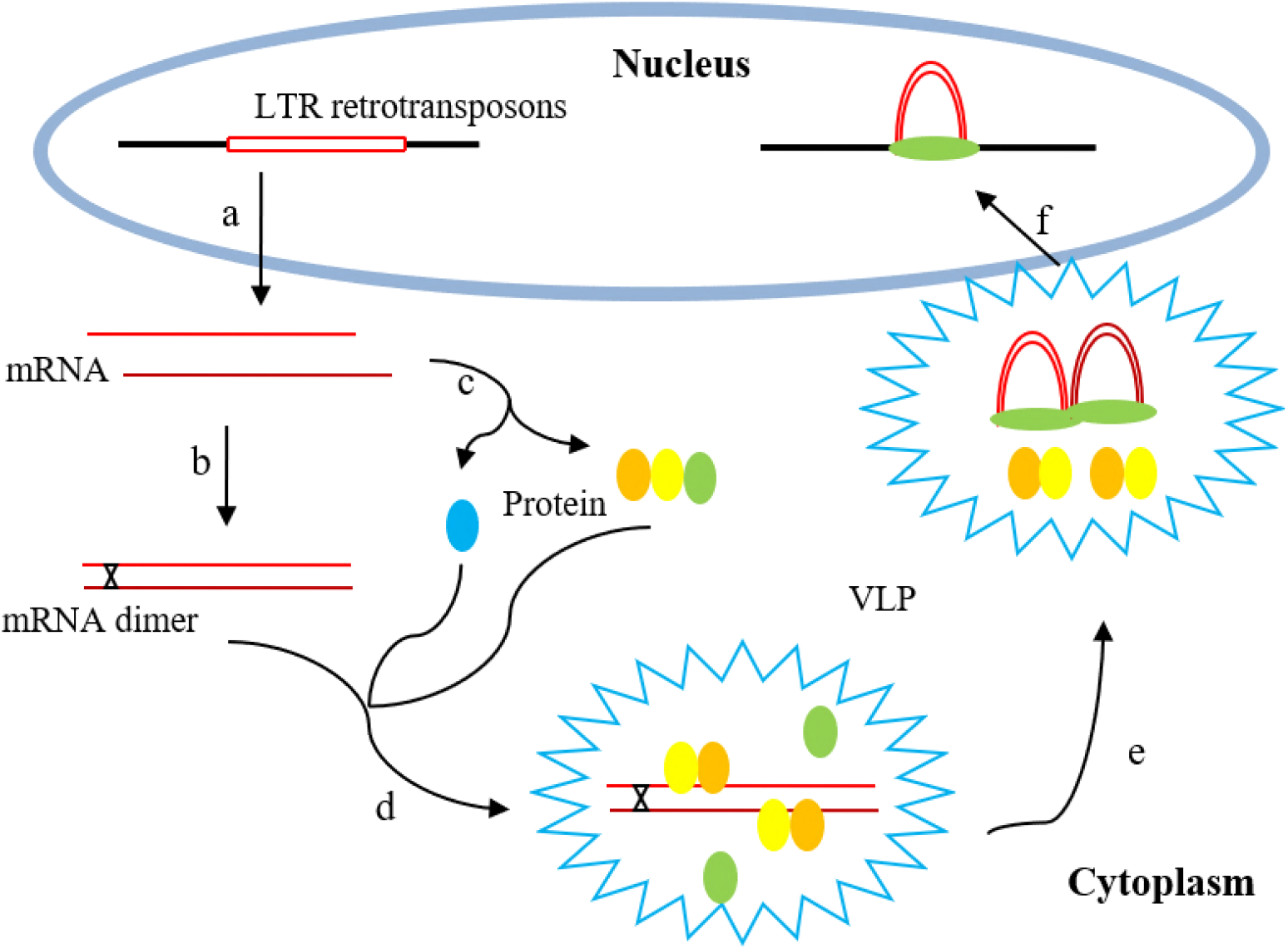
The life cycle of LTR retrotransposons. (a) LTR retrotransposons transcribe to mRNA from host genome. (b) The dimer is generated from mRNA. (c) The mRNA translates to GAG and POL (AP, RT-RNAseH, IN). (d) The virus-like particle is formed by the GAGs polymerizing. Meanwhile, the dimer and POL proteins are packaged into the virus-like particle. (e) The double-stranded cDNA is formed by reverse transcription. Then the virus-like particle is localized to the nucleus. (f) LTR retrotransposons nest into the host genome.

Previously, two models had been proposed to account for the origin of the complex retrotransposons. But they differ in how and when the retrotransposons were presumed to create. Devos (Devos *et al*. 2002) had proposed that unequal intra-strand homologous recombination between LTRs of different normal elements that belonging to the same family can create the complex LTR retrotransposons, which was the result of genome structural variation and occurred after inserting the host genome. While, Sabot (Sabot and Schulman 2007) suggested another model that the complex LTR retrotransposon was inserted by an abnormal template-switching during the reverse transcription process and it had formed a new copy before inserting into the host genome. Both models have important implications for exploring the origins and evolution of complex LTR retrotransposons. But they can’t account for the origin of these complex elements in which outermost LTRs do not harbor the most similarities. Here, we will describe a new mechanism to account for the complex LTR retrotransposons formation before their integration to the host genome.

The genus *Oryza* was an ideal model system to study formation of the complex LTR retrotransposons. *Oryza sativa* genome was sequenced and further effort still continued in order to improve the genome sequence quality (Matsumoto *et al*. 2005). Furthermore, there have been a few detailed studies of LTR retrotransposons in the rice genome (McCarthy *et al*. 2002; Ma and Bennetzen 2004; Ma *et al*. 2004; Vitte *et al*. 2007; Zuccolo *et al*. 2007; Takuno and Gaut 2012). Previous study indicate that LTR retrotransposons are the major component of the rice genome (Zuccolo *et al*. 2007), which have a rapidly recent growth and follow with a rapidly genomic DNA loss (Ma and Bennetzen 2004; Ma *et al*. 2004). Many young normal LTR retrotransposons were created in rice genome by recent amplification.

## Results and Discussion

Using LTRtype based on Repbase library (Jurka *et al*. 2005) (Additional file 1) and further carefully manual inspection, we clearly identified 63 complex LTR retrotransposons in the rice genome (Table 1). A careful analysis of TSDs of available 63 complex elements showed that 45 complex elements harbored TSDs. It suggested that the origin of two outermost LTRs (LTR1, LTR3) belonging to the single elements of the forty-five were from a single insertion event. Moreover, another 18 complex elements did not harbor flanking TSDs, which suggested that they were derived from unequal intra-strand homologous recombination in host genome. This is consistent with Devos’ study (Devos *et al*. 2002). However, it can’t account for other forty-five elements.

**Table 1:**
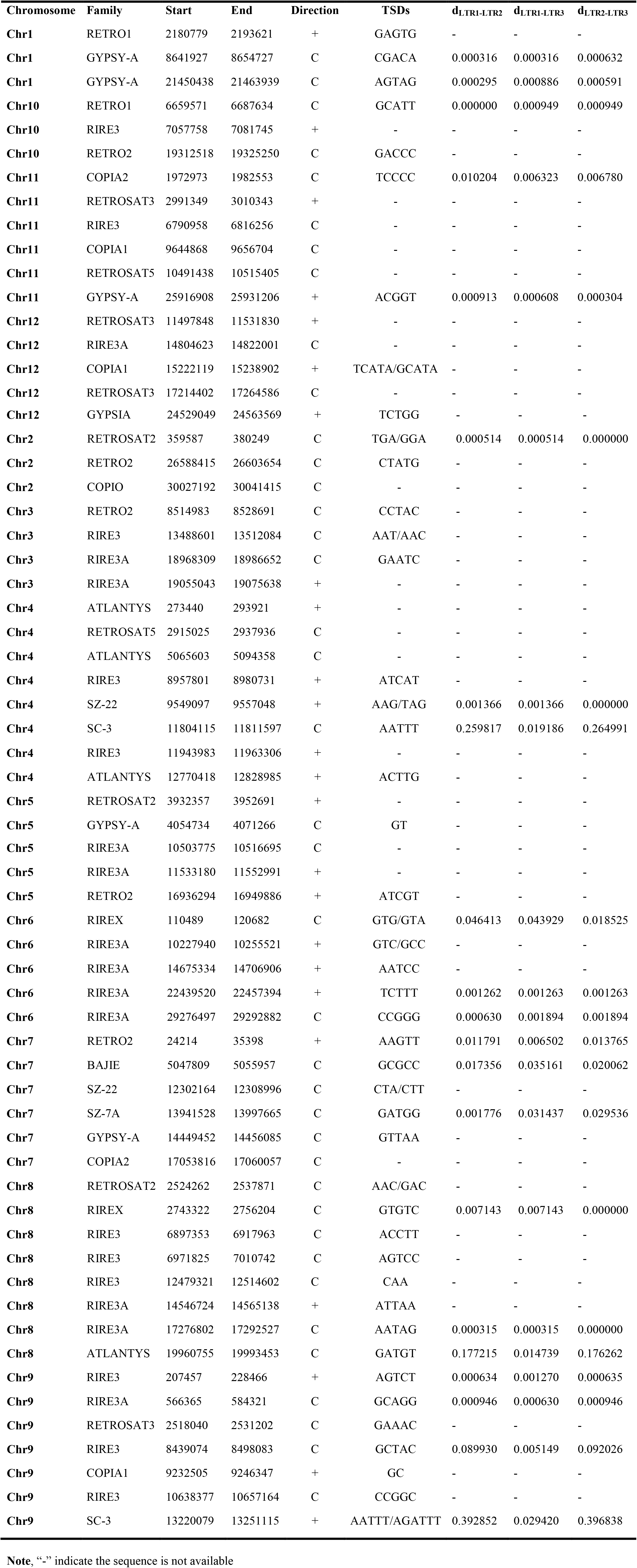
The information of the complex LTR retrotransposons in rice genome.

Among the 45 complex elements, twenty-one complex elements’ LTRs were not completely lost. These incomplete elements are excluded in the later study. Then, the genetic distances of three LTRs (LTR1—LTR2—LTR3) were calculated in each complex LTR retrotransposons of these completely elements (Table 1). Their genetic distances were expressed as d_LTR1-LTR2_, d_LTR1-LTR3_, and d_LTR2-LTR3_, respectively. The genetic distances of two flanking LTRs (d_LTR1-LTR3_) were the smallest in six elements. It indicated that LTR1 and LTR3 were the most highly homologous. These complex elements can be explained by Sabot’s study (Sabot and Schulman 2007). However, some complex elements had different characteristics. d_LTR1-LTR3_ was equal to d_LTR1-LTR2_ or d_LTR2-LTR3_ in eight complex LTR retrotransposons. Among them, five elements in which d_LTR1-LTR3_ equaled d_LTR1-LTR2_ suggested that LTR2 and LTR3 in the single element were from a single integration event. Other three elements in which d_LTR1-LTR3_ equaled d_LTR2-LTR3_ suggested that LTR1 and LTR2 in the single element were from a single integration event. Combined with the result of the previous paragraph, we concluded that three LTRs (LTR1—LTR2—LTR3) of the single complex element were from a single integration event in all eight complex elements. Moreover, in three cases, d_LTR1-LTR3_ approximately equaled d_LTR1-LTR2_ or d_LTR2-LTR3,_ suggesting that it had the same origin with the front eight elements. Although Sabot proposes that the abnormal template switching can create complex LTR retrotransposons insertion in genome which result in their two flanking LTRs of the set of three being highly homologous (Sabot and Schulman 2007), it still can’t explain eleven complex retrotransposon elements.

We thus suggest a new model that the complete LTRs homologous recombination of two different normal LTR retrotransposon elements of the same family before inserting the host genome which can explain eleven complex retrotransposon elements formation (Figure 2.A.a1). Two normal LTR retrotransposon elements are formed from two different normal LTR retrotransposons of the same family in host genome via transcription and reverse transcription. The LTRs are the same in each new element. Because of their LTRs having homologous in two elements of the same family, they can occur recombination, and then form complex LTR retrotransposon elements harbored three LTRs. The new complex LTR retrotransposon inserts in host genome and produce flanking TSDs. An important characteristic of the new insertion complex retrotransposon is that one of the outermost LTRs (LTR1 or LTR3) has the same genetic distance to the other two LTRs (LTR2, LTR3 or LTR1).

**Figure 2:**
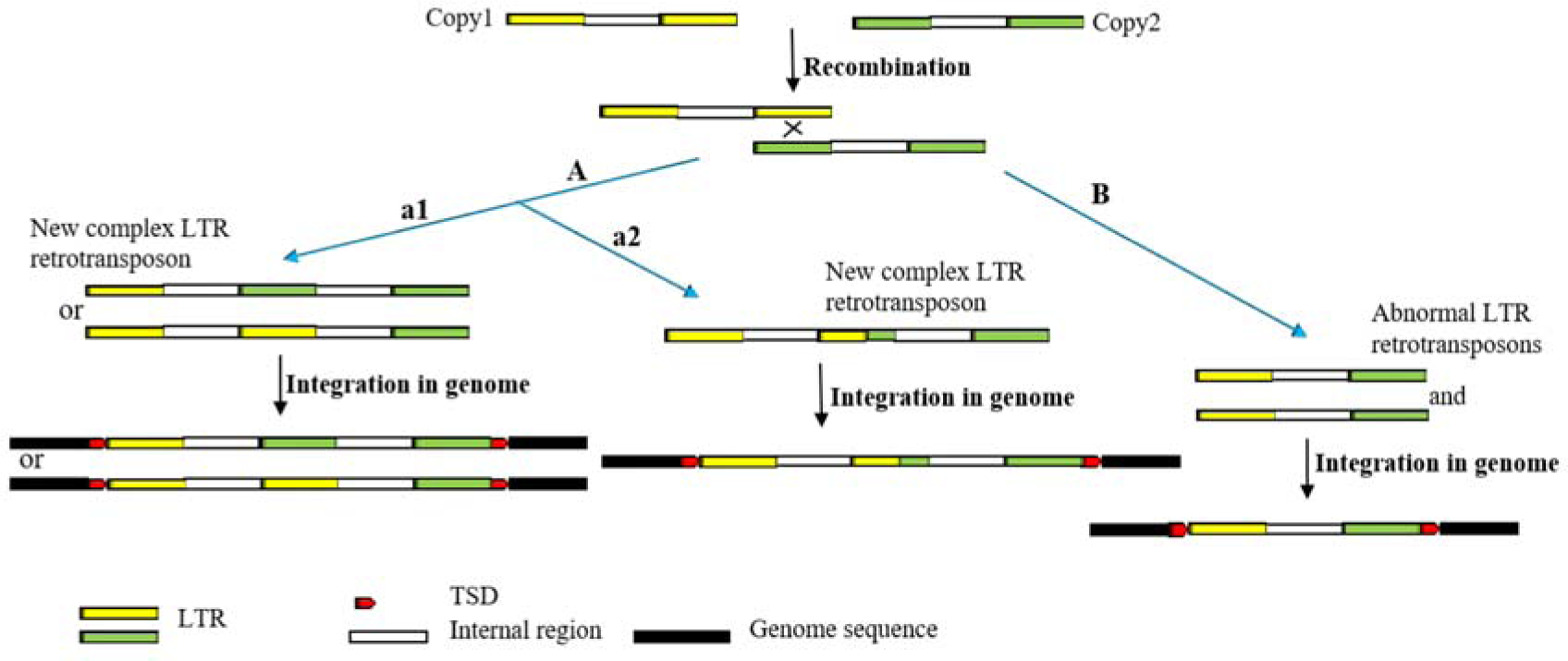
Two new copies have been synthesized from different normal LTR retrotransposons of the same family by reverse transcription. In the highly homologous LTRs region, two copies can be occurred recombination. A: a complex LTR retrotransposon and a solo LTR are formed. Then the complex LTR retrotransposons insert in host genome resulting in harbors flanking TSDs. B: two abnormal LTR retrotransposons are formed, and two LTRs are different in each element. Then they insert in host genome and result in harbors flanking TSDs.

Retrotransposons are thought to share many features with the life cycle of retroviruses (Sabot and Schulman 2006) (Figure 1). The retroviruses RNA is generally dimeric within the virus-like particle (VLP), and it either occurs before or simultaneously with packaging (Brunel *et al*. 2002). A LTR retrotransposon element had been shown to be dimeric in the bakers’ yeast *Saccharomyces cerevisiae* (Feng *et al*. 2000). The double-stranded cDNA was synthesized in the VLP. The VLP was ultimately localized to the nucleus and the double-stranded cDNA transferred into the nucleus. In this life process, dimerization and VLP provided the source and place for homologous recombination of retrotransposons before their integration, respectively. The tools of homologous recombination were generated by missing packaging. In the packaging process of RNA and polyprotein, we thought that recombinant proteins were packaged into the VLP. Before this, there were many studies for homologous recombination of retrovirus (Hu and Temin 1990; Stuhlmann and Berg 1992; Taucher *et al*. 2010; Delviks-Frankenberry *et al*. 2013). The two models had been proposed to understand it. The first model was proposed by Coffin (Coffin 1979), it was a modified copy choice mechanism in which reverse transcriptase switches from one RNA template to another upon encountering breaks in the RNA stands. Abnormal template switching in the DNA negative-strand synthesis of reverse transcription was a similar mechanism with Sabot’s study in plant retrotransposons (Sabot and Schulman 2007). Another model was that the two RNA genomes were each reverse transcribed into negative-strand DNA and that single-stranded DNA branches were formed and recombine with homologous regions on the other cDNA in a displacement-assimilation mechanism (Junghans *et al*. 1982). In recent year, recombination of retrotransposons and exogenous RNA virus had been identified in mammal (Geuking *et al*. 2009).

To further validate this model, we obtained 249 normal LTR retrotransposons from the RIREX family using the same method as before (Additional file 2). RIREX was a large family and contains two complex LTR retrotransposons in rice genome. These normal elements were named as 1~249. The LTRs of all normal elements were analyzed for their phylogenetic relationships. This result showed that most of the single normal element‘ flanking LTRs had the closest phylogenetic relationship. However, they were especially in four normal elements, the phylogenetic relationship of single normal element’ flanking LTRs were not close relative (Figure 3). That was to say, in the four normal elements, the flanking LTRs of single normal element were from different ancestors. Further, a careful analysis of TSDs of the four normal elements showed that only one element did not harbor flanking TSDs. The above results are consistent with Devos’ study (Devos *et al*. 2002). The origin of the one element was that unequal intra-strand homologous recombination between LTRs of different elements belonging to the same family in genome. However, in other three normal LTR retrotransposons, the flanking LTRs of single element came from the homologous recombination of LTRs of different elements belonging to the same family before inserting the host genome (Figure 2.B).

**Figure 3:**
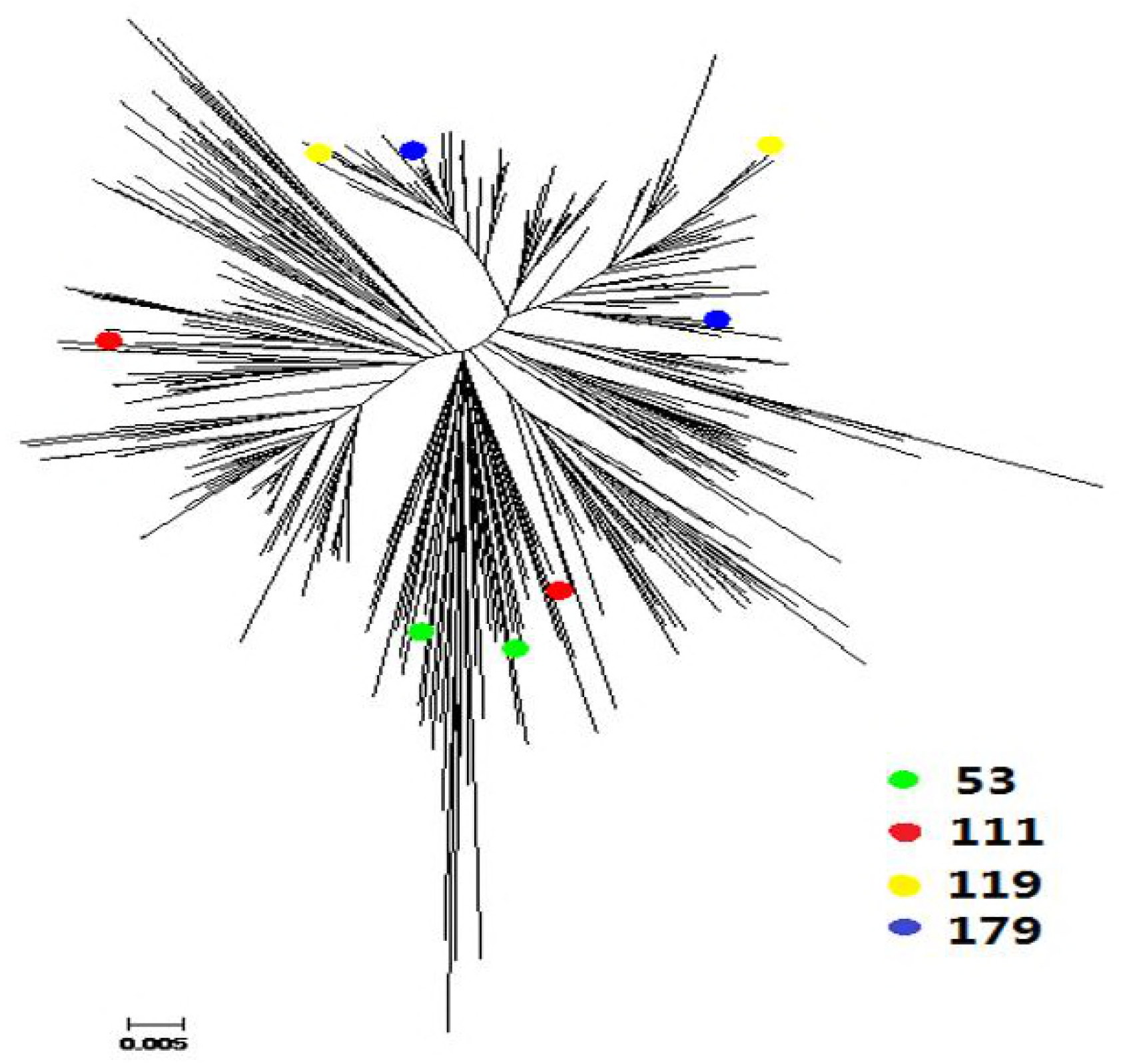
Phylogenetic analysis of all normal elements’ LTRs of RIREX family. Solid green circle represents the element 53, solid red circle represents the element 111 (TSDs: TTACC), solid yellow circle represents the element 119 (TSDs: GAC/GAA), solid blue circle represents the element 179 (TSDs: CATAG)

Furthermore, the formation of four complete complex elements is still unclear. Their LTRs sequences were analyzed by Clustalw (Thompson *et al*. 2002). In three LTRs of the complex BAJIE (Chr7, 5047809 ~ 5055957), the previous section of LTR2 was the same with LTR3, but the rest part was highly similar with LTR1 (Additional file 3). The result showed that the origin of LTR2 was from LTR1 and LTR3. The similar mechanism with above eleven complex elements that partial LTRs homologous recombination of two new different normal LTR retrotransposon copies of the same family before inserting the host genome can understand the complex element (Figure 2.A.a2). Further, two complex GYPSY-A (Chr1, 21450438 ~ 21463939; Chr11, 25916908 ~ 25931206) elements had a similar characteristic with above 12 complex LTR retrotransposons (Additional file 4, 5). They had occurred homologous recombination before their integration, but we could not identify that complete or partial LTRs occurred homologous recombination. Regrettably, although these LTRs sequence had been analyzed (Additional file 6), the origin of RIRE3 (Chr9, 207457 ~ 228466) was still a secret.

## Conclusions

Among the 63 complex elements we observe in rice, we can explain the origin of fourteen complex elements by the new model. Although the new mechanism that our study proposed can understand the formation of complex LTR retrotransposons, the origin and evolution of the complex elements are still not completely clear. Further research need to be done on the complex LTR retrotransposons.

## Methods

### Mining of complex LTR retrotransposons

The rice IRGSP/RAP genome sequences (Matsumoto *et al*. 2005) (version 7.0) were downloaded from http://rice.plantbiology.msu.edu. And the LTR retrotransposon sequences of rice were downloaded from Repbase version 18.11 (Jurka *et al*. 2005) to build the LTR retrotransposon library file that including a total of 97 pair LTRs and IRs (internal regions) (Additional file1). We annotated the rice genome by using LTRtype for complex LTR retrotransposons mining (Zeng *et al*. 2017).

### LTRs sequence analysis

The LTRs sequences were aligned with Clustalw (Thompson *et al*. 2002). Then, the genetics distances of both LTRs were estimation using the Kimura two parameters method (Kimura 1980), calculated using MEGA5 (Tamura *et al*. 2011). The phylogenetic relationships of LTRs of normal full length LTR retrotransposons were estimation using NJ method, implemented in MEGA5 (Tamura *et al*. 2011).

### Additional file

Additional file 1 is the library of LTR retrotransposons from Repbase; and Additional file 2 is these LTRs sequence from 249 normal elements of RIREX family. Additional file 3, 4, 5 and 6 are the alignment sequence about four complex LTR retrotransposons by Clustalw.

## Acknowledgements

The authors would like to thank LiZhi Gao, Shuo Wang and Chao Shi for their technical support and infinite helpfulness.

## Abbreviations

LTR: Long terminal repeat; TSDs: target-site duplications; ORF: open reading frame; VLP: virus-like particle

## Availability of data and materials

Not applicable.

## Authors’ contributions

Coordinated and drafted the manuscript: ZY Sun, FC Zeng contributed to Figures and table. ZY Sun and FC Zeng authors contributed to manuscript writing and read and approved the final version of this article

## Consent for publication

Not applicable.

## Ethics approval and consent to participate

Not applicable

